# Deep transcriptome profiling of human hypothalamic agouti-related protein and proopiomelanocortin neurons regulating energy homeostasis

**DOI:** 10.1101/2025.08.19.671088

**Authors:** Szabolcs Takács, Katalin Skrapits, Balázs Göcz, Éva Rumpler, Miklós Sárvári, Szilárd Póliska, Gergely Rácz, András Matolcsy, Erik Hrabovszky

## Abstract

We have developed and validated a pioneering ‘IHC/LCM-Seq’ method for transcriptome profiling of spatially defined neuronal cell types detected with immunohistochemistry in sections of formaldehyde-fixed human brains. IHC/LCM-Seq provided unprecedented insight, with 14,000-16,000 transcripts identified, into the gene expression landscape of human agouti-related peptide (AgRP) neurons, which drive appetite and energy storage, and proopiomelanocortin (POMC) neurons, which suppress feeding and promote energy expenditure. These cells differed from each other, and from fertility-regulating kisspeptin neurons, in their distinct enrichments of co-transmitters, transcription factors, and receptors. The AgRP neuron transcriptome was rich in receptors for proinflammatory cytokines, metabolic hormones and growth hormone, whereas POMC neurons expressed reproductive hormone-, glucagon-like peptide– and endocannabinoid receptors. IHC/LCM-Seq, a versatile spatial transcriptomic approach for characterizing cell types in *postmortem* brains, opened a new window onto molecular mechanisms regulating energy homeostasis in the human hypothalamus and highlighted possible pharmacological targets for weight management.

## Introduction

The hypothalamic arcuate (aka infundibular) nucleus plays a crucial role in energy homeostasis, primarily through the actions of two functionally antagonistic neuronal populations: Agouti-related protein (AgRP) neurons, which stimulate appetite and promote energy conservation, and proopiomelanocortin (POMC) neurons, which suppress food intake and enhance energy expenditure.^1^ AgRP and POMC neurons are highly responsive to peripheral metabolic cues – including leptin, insulin, and ghrelin – and their dysfunction is strongly linked to disorders of energy homeostasis, such as obesity, anorexia, and diabetes.^1,2^ Defining the molecular signatures of AgRP and POMC neurons is fundamental for identifying therapeutic strategies against metabolic disorders. In particular, detailed characterization of their receptor profiles may uncover novel mechanisms regulating energy homeostasis and guide targeted approaches to address the obesity pandemic. For example, expression of *GLP1R* in anorexigenic POMC neurons likely contributes to the anti-obesity effects of semaglutide, whereas calcitonin receptor (*CALCR*) expression in these neurons may underlie the efficacy of recent combination therapies pairing semaglutide with the amylin analogue cagrilintide, which can act on CALCR.^3^

Current knowledge of the regulatory mechanisms within hypothalamic neuronal circuits targeted by obesity and diabetes treatments has largely been derived from animal studies, limiting the translational potential for developing effective therapies in humans. Recent advances in single-nucleus RNA sequencing (snRNA-seq) have enabled the creation of a spatio-cellular transcriptional map (‘HYPOMAP’) of the human hypothalamus.^4^ The authors identified approximately 2,000 transcripts per cell and integrated their snRNA-seq data with a hypothalamic expression matrix derived from publicly available whole-brain human datasets.^5^ Through multi-level hierarchical unsupervised clustering, they delineated 291 hypothalamic neuronal clusters, including a single AgRP neuron cluster defined by five signature genes, and three distinct POMC neuron clusters characterized by eight marker genes.^4^

The small quantity of input RNA severely limits the sensitivity of various single-cell sequencing techniques. By contrast, bulk sequencing of neuronal pools isolated via fluorescence-activated cell sorting (FACS) or laser capture microdissection (LCM) enables the detection of rare transcripts and produces highly detailed transcriptomic profiles that are otherwise difficult to obtain. Our laboratory recently achieved a methodological breakthrough in bulk sequencing by integrating innovative RNA-preserving strategies during immunohistochemistry (IHC) with the isolation of labeled neurons using LCM, followed by RNA-Seq.^6^ In the present study, we adapted the ‘IHC/LCM-Seq’ approach to human *postmortem* brains fixed by immersion in 4% formaldehyde (FA). This adaptation enabled transcriptomic profiling of AgRP and POMC neurons in the arcuate/infundibular nucleus, as well as fertility-regulating kisspeptin (KP) neurons^7^ of the same region, which we selected for comparison with a third cell type. The IHC/LCM-Seq methodology yielded unprecedented transcriptome coverage – approximately 14,000 to 16,000 transcripts per cell type. These included rare transcription factors, regulatory RNAs, and, notably, receptors, some of which may become potential targets for future weight-management strategies.

## Results

### Immersion-fixation of *postmortem* brain samples preserves tissue RNA integrity

Perfusion fixation of the brain with 4% FA constituted the initial step of the IHC/LCM-Seq protocol we developed for laboratory rodents.^6^ Because this fixation protocol is not applicable to autopsy material, we first assessed how *postmortem* delay and immersion fixation jointly affect tissue RNA integrity. Neocortical blocks (∼4 mm in each dimension) were dissected from the brain of a 53-year-old woman ∼20 h after death and immersion-fixed in cold (4 °C) 4% FA for 24, 48, or 72 h, then infiltrated with 20% sucrose for 24 h. Twenty-µm-thick free-floating sections were prepared using a freezing microtome and stored at – 20 °C in a cryoprotectant solution formulated for long-term RNA preservation^6^ (**Fig. 1A**). As we proposed recently^6^, all solutions included the RNase inhibitor aluminon^8^ (aurintricarboxylic acid, ammonium salt; ATA; 0.05%). Bioanalyzer measurements of total RNA extracted from test sections revealed consistently high RNA integrity (RIN > 7.0) and comparable yields (∼2 ng/mm^2^), regardless of fixation duration (**Fig. 1B**, **Table S1**).

**Fig. 1:**
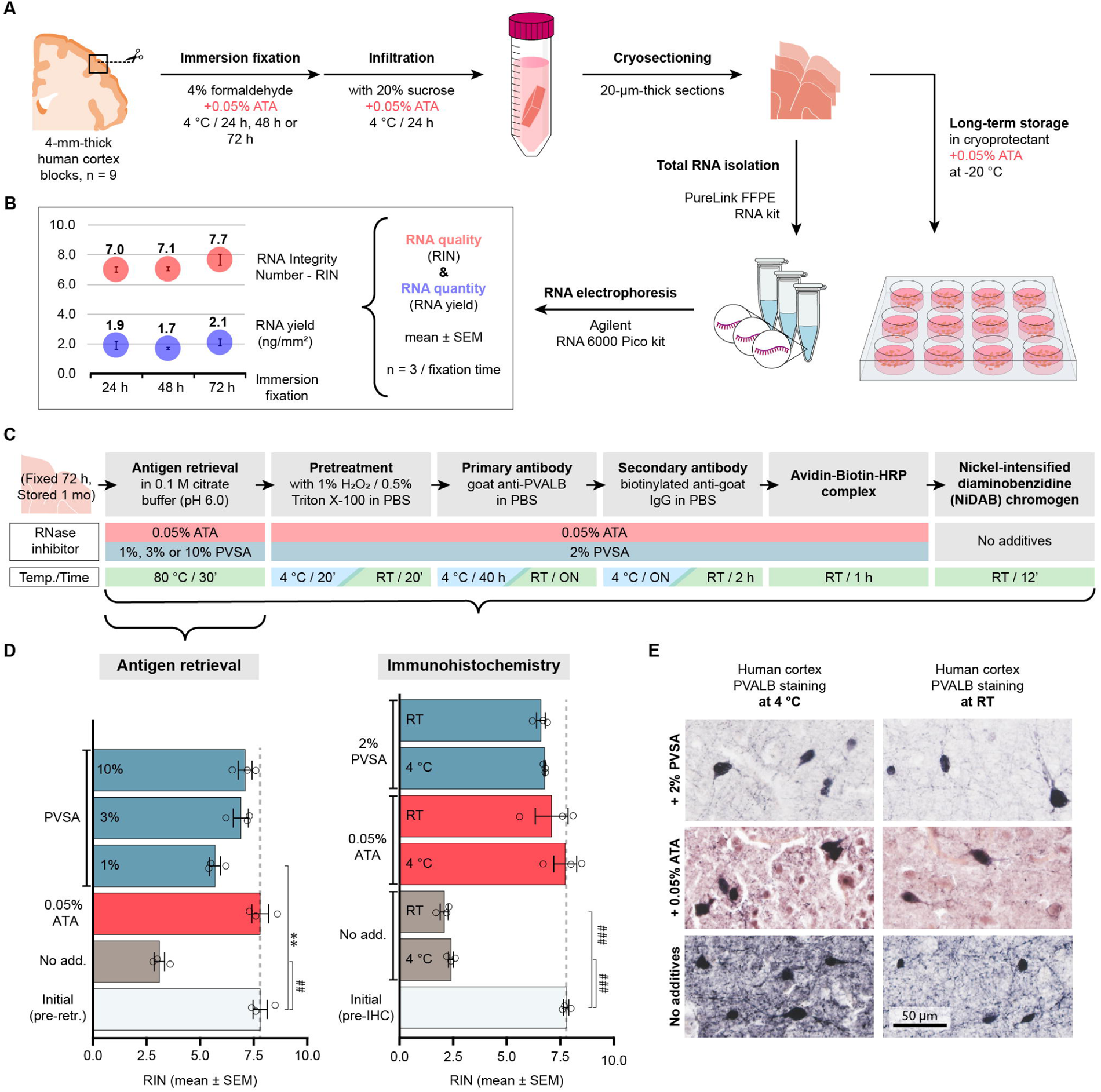
Development and validation of a versatile ‘IHC/LCM-Seq’ method for transcriptome profiling of immunohistochemically-labeled human neurons. **A**: Schematic overview of histological procedures used to evaluate the combined effects of *postmortem* delay and immersion fixation (24, 48, or 72 h) on RNA integrity. **B**: Neocortical tissue samples from a 53-year-old female, processed in the presence of the RNase inhibitor aluminon (ATA; 0.05%), show comparable RNA quality (RIN > 7.0) and yield across different fixation times (Agilent Bioanalyzer; Table S1). **C**: Experimental design for optimizing RNA preservation using ATA (0.05%) or polyvinyl sulfonic acid (PVSA; 1-10%) during immunohistochemical detection of cortical parvalbumin (PVALB) neurons. **D**: Both ATA and PVSA provide significant protection against degradation during antigen retrieval at 80 °C, while RNA integrity is severely compromised in the absence of RNase inhibitors; ATA and PVSA (2%) also maintain RNA integrity during IHC performed either at RT or 4 °C (Table S1). **E**: Effects of ATA (0.05%) and PVSA (2%) on IHC signal quality and background. Both additives enable successful detection of PVALB neurons; however, ATA treatment results in increased nonspecific background staining. Statistical annotations: **p<0.01, ##p<0.00001, ###p<0.000001 by one-way ANOVA followed by Tukey’s HSD.

### A modified immunohistochemistry protocol preserves tissue RNAs

Encouraged by the high RIN values of immersion fixed samples, we next adapted the IHC/LCM-Seq workflow to human tissue sections. The modified IHC protocol began with epitope retrieval in 0.1 M citrate buffer (pH 6.0, 80 °C, 30 min), supplemented with either 0.05% ATA or polyvinyl sulfonate (PVSA) at concentrations of 1%, 3%, or 10% (**Fig. 1C**). In the absence of RNase inhibitors, RIN values dropped sharply from 7.8 ± 0.3 to 3.1 ± 0.2. In contrast, inclusion of 0.05% ATA or 3-10% PVSA effectively preserved RNA integrity (**Fig. 1D, Table S1**). As a final validation of our RNA-preserving IHC protocol, we visualized cortical parvalbumin neurons immunohistochemically (**Fig. 1E**), followed by RNA quality analysis. Sections immunostained without RNase inhibitors exhibited substantial RNA degradation, with RIN values of 2.4 ± 0.1 at 4 °C and 2.1 ± 0.2 at RT. In contrast, supplementing all antibody and buffer solutions with 0.05% ATA or 2% PVSA effectively preserved RNA, yielding RIN values of 7.7 ± 0.5 (ATA) and 6.8 ± 0.0 (PVSA) at 4 °C, and 7.1 ± 0.8 (ATA) and 6.6 ± 0.2 (PVSA) at RT (**Fig. 1D**, **Table S1**).

Of note, ATA tended to compromise specific immunostaining and increase nonspecific background (**Fig. 1E**), consistent with our previous observations.^6^ Therefore, the finalized IHC/LCM-Seq protocol for profiling hypothalamic AgRP, POMC, and KP neurons incorporated 2% PVSA in all buffer and antibody solutions (**Fig. 2A**).

**Fig. 2:**
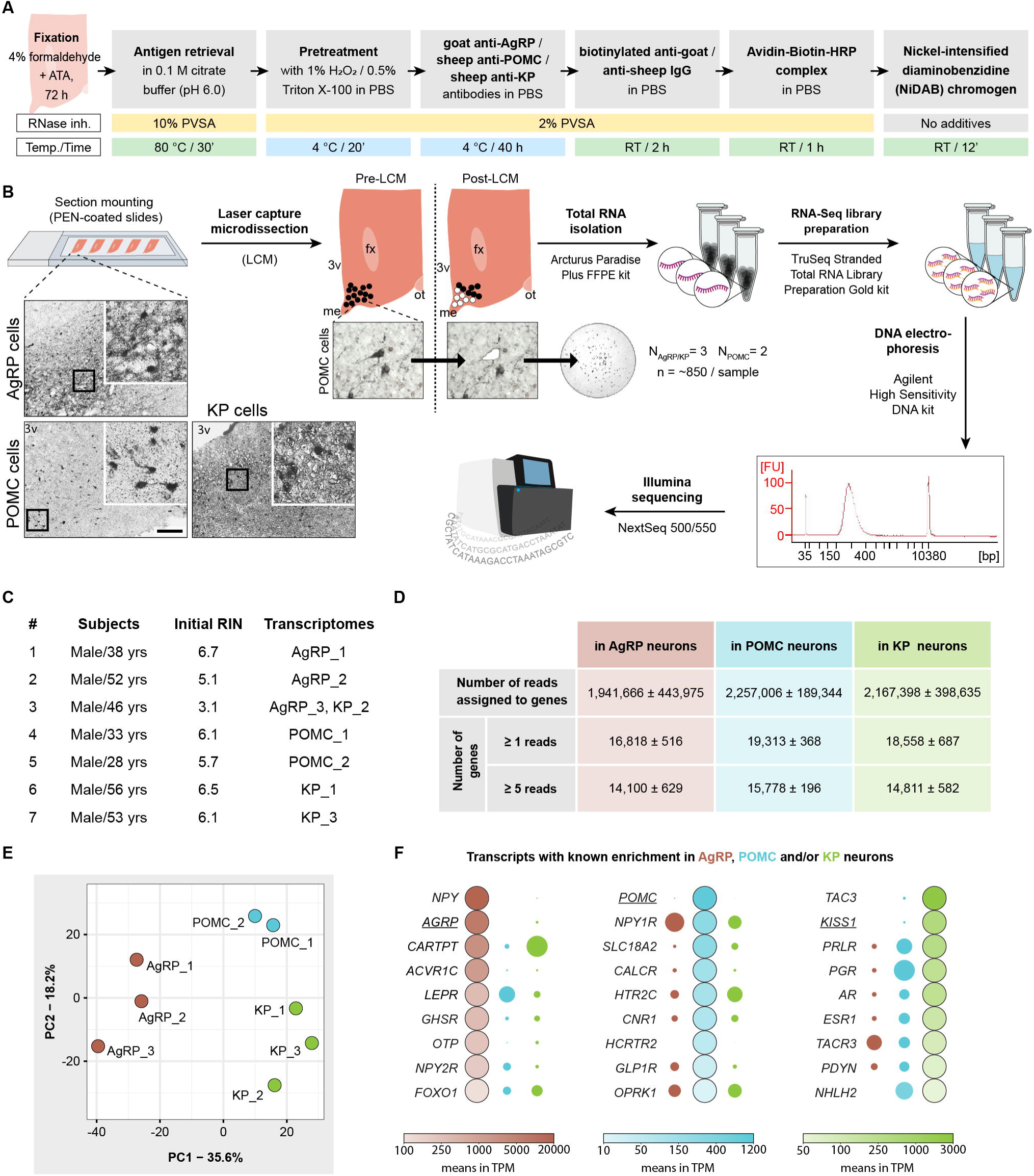
Use of IHC/LCM-Seq for transcriptome profiling of human AgRP, POMC and KP neurons. **A**: Schematic overview and technical details of the RNA-preserving immunohistochemistry (IHC) protocol optimized for transcriptome analysis of human AgRP-, POMC-, and KP-immunoreactive neurons. **B:** Workflow of the protocol, including laser capture microdissection (LCM), RNA isolation, cDNA library preparation, RNA sequencing, and downstream bioinformatic analyses. Scale bar represents 200 µm in overview images and 50 µm in high-magnification LCM images. 3v – third ventricle, fx – fornix, me – median eminence, ot – optic tract. **C:** Summary of donor and tissue parameters for samples used in transcriptomic profiling of the three neuronal populations. **D:** Number of gene-assigned reads and expressed genes identified in AgRP, POMC, and KP neurons at two expression thresholds (**Table S2**). **E:** Principal component analysis (PCA) showing distinct clustering of AgRP, POMC, and KP transcriptomes, with PC1 and PC2 explaining ∼50% of the total variance. **F:** Representative genes enriched in a cell type-specific manner. Color gradients within the large dots indicate transcript abundance in mean TPM (transcripts per million), while the surface area of the smaller dots is proportionally reduced to reflect lower expression levels.

### Optimized tissue processing enables high-resolution transcriptome profiling of hypothalamic AgRP, POMC and KP neurons

The optimized IHC/LCM-Seq protocol (**Fig. 2A**) enabled robust immunohistochemical detection of AgRP, POMC and KP neurons in the human hypothalamus, facilitating their capture by LCM (∼850 labeled cells per cell type per subject). This was followed by RNA isolation, cDNA library preparation, and Illumina-based RNA sequencing (**Fig. 2B**). Pre-IHC RINs from hypothalamic test sections of seven male subjects ranged from 3.1 to 6.7 (**Fig. 2C**). Sequencing data have been deposited in BioProject under accession numbers PRJNA1305226 (AgRP and POMC neurons) and PRJNA1305334 (KP neurons).

Sequencing of eight libraries – comprising 3 AgRP, 2 POMC, and 3 KP samples – generated a total of 211.8 million raw reads (17.6-33 M per sample). Following trimming, adapter removal, alignment to the human reference genome (Ensembl release 111) using STAR (v2.7.11b), and gene assignment with FeatureCounts (subread v2.0.6), an average of 2.2 ± 0.6 million reads per sample (17.6 million total) were uniquely mapped and assigned to genes. This enabled identification of approximately 18,000 ± 1200 transcripts with ≥ 1 read, and 15,000 ± 800 transcripts with ≥ 5 reads (**Fig. 2D**, **Table S2**). Principal component analysis (PCA) verified three distinct transcriptomic profiles, each corresponding to a specific cell type (**Fig. 2E**). The absence of marker neuropeptide transcripts (*AGRP*, *POMC*, and *KISS1*) outside their respective neuronal populations, together with asymmetric expression of established neuronal phenotype markers, confirmed that there was minimal – if any – cross-contamination between transcriptomes **(Fig. 2F)**.

### Differential gene expression analysis reveals cell type-specific transcript enrichment, including 50 transcription factors

A total of 1,210 transcripts were significantly enriched in a cell type-specific manner among AgRP, POMC, and KP neurons (adjusted p-value /p.adj./ < 0.05, DESeq2; **Table S2**). Differentially expressed genes (DEGs) are visualized through heat maps (**Fig. 3A**), numerical expression data (**Fig. 3B**), and volcano plot markers across the three pairwise comparisons (**Figs. 3C-E**). We further examined 902 transcription factor (TF) transcripts annotated in the KEGG BRITE database to explore their potential roles in shaping the distinct transcriptomic profiles of AgRP, POMC, and KP neurons (**Table S3**). Of the 40 most abundant TFs selected per cell type (63 in total), 13 were significantly differentially expressed, indicating cell type-specific transcriptional regulation (**Fig. 3F**, **Table S3**). Expanding the analysis to all expressed TFs revealed 50 that differed between at least two neuronal populations, underscoring broader cell type-specific transcriptional regulation (p.adj. < 0.05, DESeq2; **Fig. 3G**, **Table S3**). Many of the shared transcription factors could reflect similar developmental programs, humoral signals, and afferent inputs. Transcription factors with significant cell type-specific enrichment (p.adj. < 0.05, fold change > 1.5 *vs.* other cell types) included *NR3C1, OTP, XBP1, KLF9, FOXO1, ETV5, CREM, ZBTB16,* and *ST18* for AgRP neurons; *ZFHX4, ONECUT1,* and *LEF1* for POMC neurons; and *BCL6, ISL1,* and *PROX1* for both AgRP and POMC neurons. Unlike AgRP neurons, POMC cells also shared expression for several transcription factors with KP neurons (*PGR, AR, ESR1, NHLH2, and SOX1*). *SOX14, STAT5A,* and *PLAGL1* were enriched in KP neurons (**Fig. 3G**; **Table S3**).

**Fig. 3.**
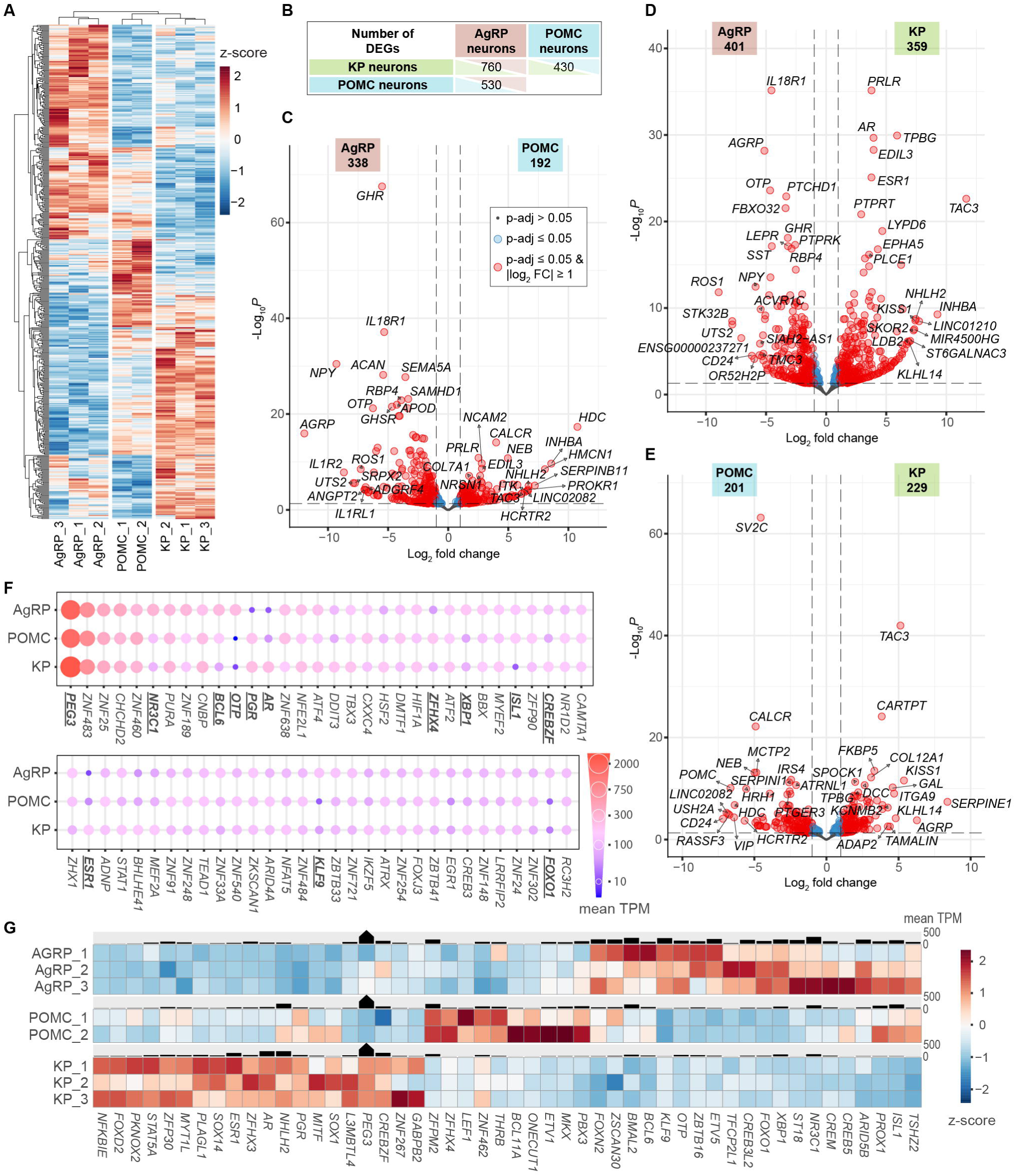
Differential expression of transcripts and transcription factors in human AgRP, POMC, and KP neurons. **A**: Heat maps showing 1,210 transcripts differentially expressed (adjusted p /p.adj./ < 0.05, DESeq2) between any two arcuate nucleus cell types. **B:** Table summarizing the number of differentially expressed genes (DEGs) in each pairwise comparison. **C-E:** Volcano plots showing the relationship between expression magnitude (log_2_ fold-change /FC/) and statistical significance (log_10_ p.adj.) across all transcripts. Gene symbols denote the top 10 upregulated and top 10 downregulated transcripts by fold change, as well as the 10 transcripts with the lowest adjusted p-values for each direction. **F:** Dot plots showing 63 transcription factors (TFs), selected as the 40 most abundantly expressed TF transcripts in each cell type. Dot size and color are proportional to expression levels (TPM). TFs with significant differential expression (p.adj. < 0.05) are marked with bold, underlined labels. **G:** Heat maps showing cell type-specific expression patterns of 50 TFs differentially expressed between AgRP, POMC and KP neurons (p.adj. < 0.05). See **Table S2** for the complete list of expressed genes and **Table S3** for expressed transcription factors.

### AgRP, POMC and KP neurons exhibit distinct neuropeptide co-transmitter profiles

Neuropeptide and granin expression profiles, based on annotations from the Neuropeptide Database^9^ (www.neuropeptides.nl), showed marked differences among the three neuronal populations. Normalized transcript abundances, expressed in TPM, are presented in **Fig. 4A** and **Table S4**. AgRP neurons were enriched for *NPY*, *AGRP*, *SST*, and *UTS2*, and exhibited higher expression of *TAC1*, *GAL*, *VGF*, and *CRH* compared to POMC and KP neurons. POMC neurons uniquely expressed *POMC* and *VIP*, and had higher levels of *ADCYAP1* relative to KP neurons. KP neurons selectively expressed *TAC3* and *KISS1*, and showed elevated expression of *CBLN1*, *NXPH1*, *PENK*, *CBLN4*, *PDYN*, and *NMU* compared to AgRP and POMC neurons (**Figs. 4A**, **B**).

**Fig. 4.**
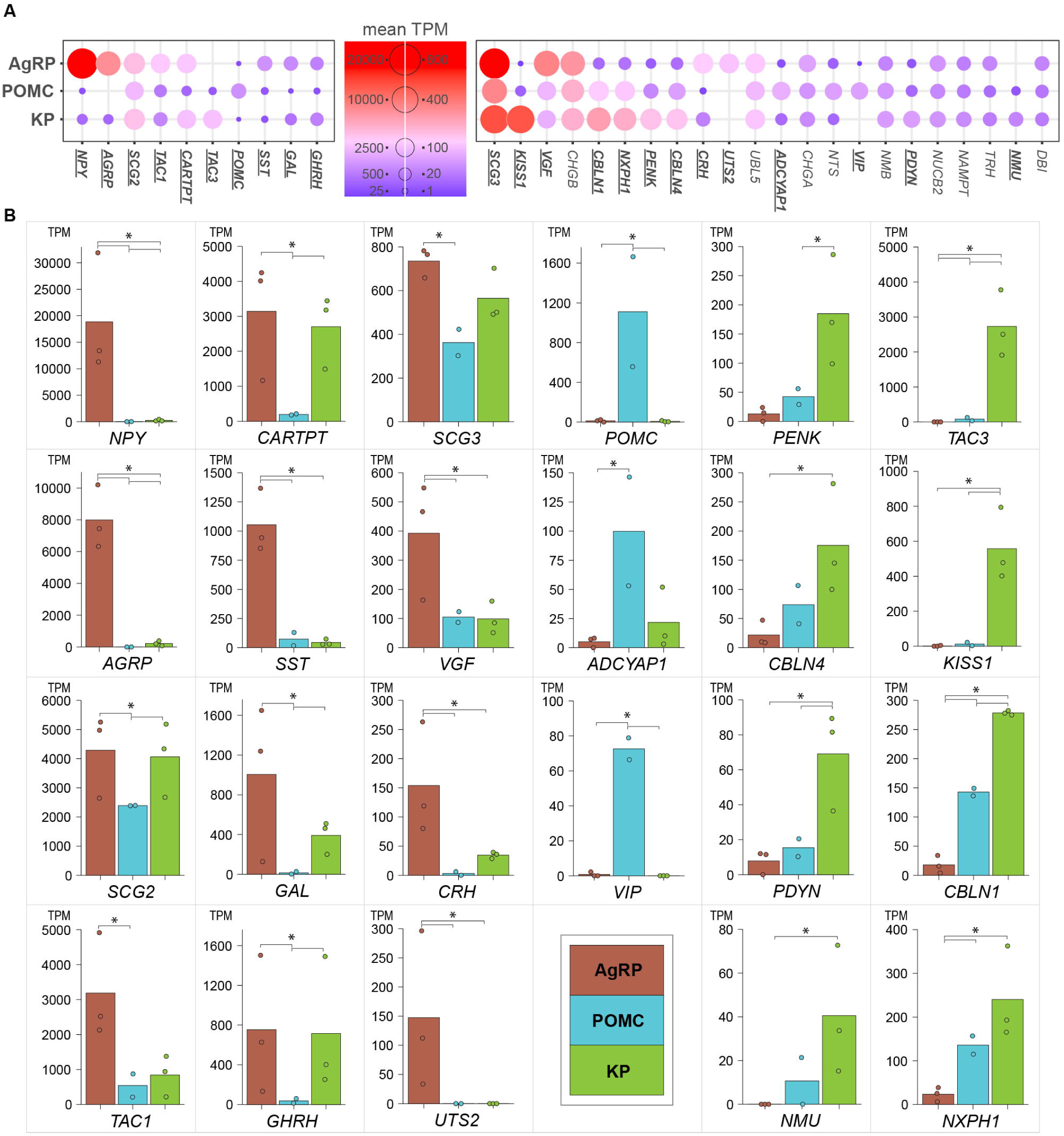
Neuropeptide co-expression in human AgRP, POMC, and KP neurons. **A**: Dot plots showing 32 neuropeptide and granin transcripts curated from the Neuropeptide Database (www.neuropeptides.nl), with mean expression levels of ≥ 40 TPM in at least one neuronal phenotype. Dot size and color are scaled according to transcript abundance. Gene symbols in bold and underlined indicate statistically significant differences between any two cell types (p.adj. < 0.05, DESeq2). **B:** Bar charts displaying the 23 neuropeptide genes that show differential expression in pairwise comparisons (*p.adj.≤ 0.05 by DEseq2). See **Table S4** for detailed expression values and statistical results for all neuropeptides.

### A subset of receptors expressed in AgRP, POMC, and/or KP neurons was implicated in genetic disorders affecting energy homeostasis

To identify transcripts associated with metabolic disorders, we selected fourteen metabolism-related traits from the GWAS Catalog^10^ (**Fig. 5A, Table S5**). These traits included 2,493 genes that were also expressed in AgRP, POMC, and/or KP neurons, applying cutoffs of mean TPM ≥ 10 and at least 5 reads in each sample of the respective cell type (**Fig. 5B, Tables S6-S7**). Among these, we identified 52 receptor genes (**Fig. 5C**, **Table S8**), several of which showed cell type-specific enrichment. Receptor expression profiles were further examined to identify potential therapeutic targets for the treatment of metabolic disorders.

**Fig. 5.**
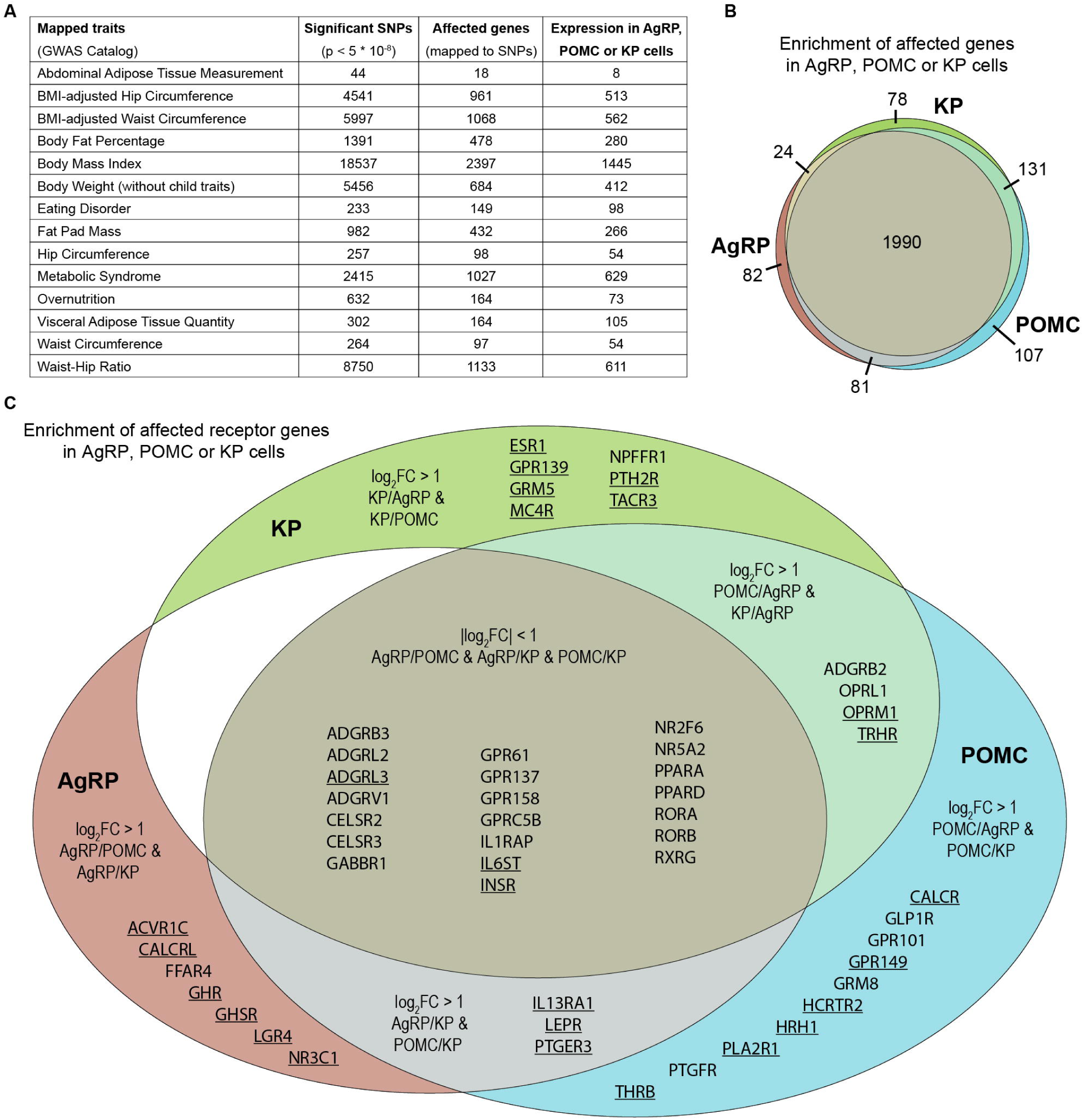
Transcripts expressed in AgRP, POMC or KP neurons that were associated with metabolic disorders. **A**: Table summarizing reported single nucleotide polymorphisms (SNPs), associated genes, and corresponding transcripts detected in AgRP, POMC, or KP neurons, linked to metabolic traits in the GWAS Catalog database (**Tables S5-S6**). Transcripts were included if they had a mean TPM ≥ 10 and at least 5 reads in each sample of the same neuronal phenotype. **B:** Cell type-dependent distribution patterns of 2,493 trait-associated transcripts (**Table S7**). **C:** Expression profiles of 52 receptor mRNAs linked to metabolic disorder traits (**Table S8**). Underlined gene symbols denote statistically significant differential expression between any two cell types (p.adj. < 0.05, DESeq2). Gene sets showing at least twofold enrichment (log_2_ fold change ≥ 1) were categorized as either cell type-specific or shared between two cell types relative to the third. (Genes in the largest intersection show no preferential enrichment in any single cell type.)

### Receptor transcripts reveal distinct humoral and neuronal regulatory mechanisms

A total of 205 receptor transcripts detectable in AgRP, POMC, and/or KP neurons (cutoffs: mean TPM ≥ 10 and reads ≥ 5 in each sample of the same cell type) are illustrated in **Fig. 6** and listed in **Table S8**. This dataset includes new candidate receptors for targeting hypothalamic metabolic and/or reproductive pathways. Of these, 112 showed at least twofold enrichment, and 62 exhibited significant differential expression in pairwise comparisons between cell types. *ACVR1C* encoding an activin receptor was the most abundantly expressed receptor, occurring exclusively in AgRP neurons. These neurons also showed selective enrichment for receptors of proinflammatory cytokines (*IL18R1*, *IL1R1*), circulating hormones (*GHR*, *GHSR*, *NR3C1*), and various neuropeptides related to energy homeostasis (*RXFP1*, *NPY2R*, *AVPR1A*, *MC3R*, *AGTR1*, and *CALCRL*). POMC neurons showed enrichment for several receptors involved in metabolic hormone and neuropeptide signaling, including *NPY1R*, *CALCR*, *SSTR1*, *HCRTR2*, *QRFPR*, *GLP1R*, *CCKAR, PROKR1,* and *GRPR*), as well as the type-1 endocannabinoid-(*CNR1*) and the H1 histamin receptor (*HRH1*). The leptin receptor (*LEPR*) was co-enriched in AgRP and POMC cells. POMC and KP neurons both expressed reproductive hormone receptors (*PRLR*, *PGR*, *AR*, *ESR1*), with higher levels in the latter, whereas opioid receptors (*OPRM1*, *OPRL1*, *OPRK1*) were expressed more prominently in POMC than in KP neurons. (**Fig. 6**; **Table S8**). *IL6ST, ADCYAP1R1*, and the insulin receptor *(INSR)* were among the most highly expressed receptors in all three cell types.

**Fig. 6.**
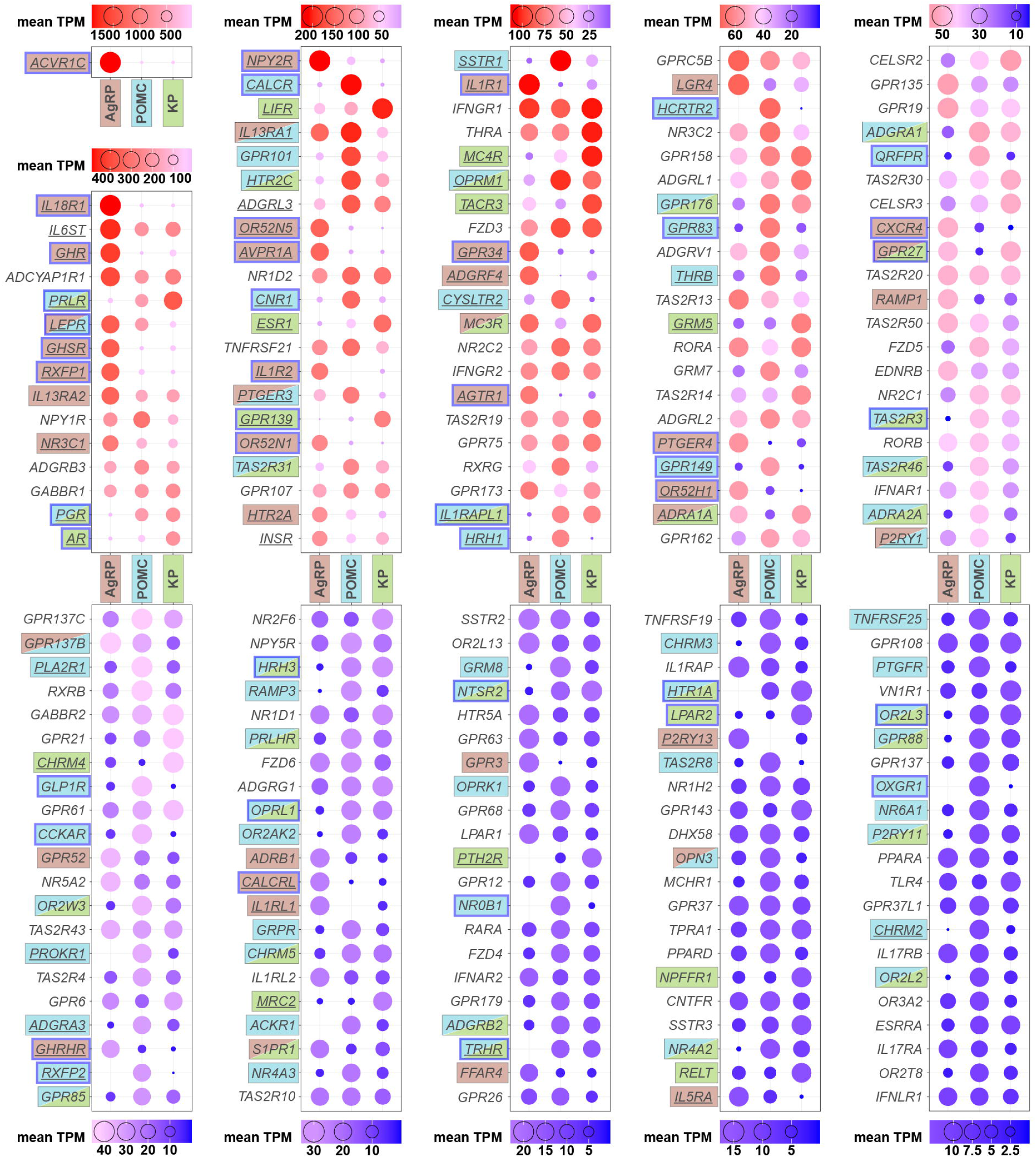
Receptor profiles of AgRP, POMC and KP neurons. Dot plots of 205 receptor transcripts (cutoffs: mean TPM ≥ 10 and reads ≥ 5 in each sample of the same cell type). Dot size and color are scaled according to transcript abundance. Underlining of symbols label statistically significant difference between any two cell types (p.adj. < 0.05, DESeq2), whereas single or double shading of the text denotes at least twofold enrichment (log_2_ fold change ≥ 1). Genes with greater than fourfold enrichment (log_2_ fold change > 2) are outlined in blue. See also **Table S8** for TPM values and statistics.

## Discussion

### Methodological considerations

The protocols used in this study are based on our previously established ‘LCM-Seq’ method, which was originally developed for spatial transcriptomic profiling of fluorescently labeled neurons in transgenic mice.^11^ To eliminate the reliance on genetic tagging, we subsequently developed the more versatile ‘IHC/LCM-Seq’ method, in which we replaced transgenic labeling with RNA-preserving immunohistochemistry. This change enabled deep transcriptomic profiling of neuronal populations across species.^6^ In the present study, we successfully adapted the IHC/LCM-Seq technique for use in *postmortem* human brain tissue. This transition introduced unique challenges inherent to autopsy-derived material, necessitating the implementation of rigorous quality control measures throughout the workflow. We found that RNA integrity was well preserved in samples collected within 24 hours *postmortem* and immersion-fixed in cold 4% FA for 24-72 hours. Variations within this fixation window had minimal – if any – impact on RNA quality, offering practical flexibility for tissue processing. Importantly, transcript detection was also largely independent of the initial RNA Integrity Number (RIN), which ranged from 3.1 to 6.7 across samples. We attributed this robustness to the use of random primers in the TruSeq library preparation protocol, which can accommodate highly fragmented RNA.^12^ Post-fixation handling – comprising cold 20% sucrose infiltration, sectioning at 20 µm, and section storage at –20 °C in cryoprotectant solution with 0.05% ATA – followed protocols we had previously optimized for mouse and rat brains.^6^ When working with immersion-fixed human brains, successful immunolabeling often requires an epitope retrieval step (30 min at 80 °C in 0.1 M citrate buffer, pH 6.0). While this pretreatment initially compromised RNA quality, the addition of 10% PVSA to the retrieval buffer effectively mitigated degradation. Subsequent immunostaining in the presence of either 0.05% ATA or 2% PVSA efficiently preserved RNA integrity at both 4 °C and RT. However, ATA caused increased nonspecific background staining and also compromised the specific signal, making PVSA the preferred RNase inhibitor for routine IHC/LCM-Seq applications. Overall, our methodological results confirmed that IHC/LCM-Seq is a robust, highly sensitive, and species-independent technique. Adapting this method to *postmortem* human brains in this study has substantial translational and clinical relevance.

As we discussed previously^6^, IHC/LCM-Seq offers several advantages over single-cell approaches such as scRNA-Seq and snRNA-Seq, which rely on enzymatic dissociation of brain tissue. i) LCM preserves the spatial context of neurons, maintaining their native anatomical architecture and microenvironment. ii) Tissue fixation prior to RNA extraction stabilizes the transcriptome and reduces artifacts introduced during dissociation and cell isolation processes. iii) IHC/LCM-Seq provides considerably enhanced sensitivity for detecting low-abundance transcripts, including long non-coding RNAs, transcription factors, and rare receptor mRNAs. However, we note that achieving unparalleled sensitivity involves a trade-off in cellular resolution. Unlike single-cell methods, bulk RNA sequencing cannot resolve cellular heterogeneity or distinguish subpopulations within a given neuronal type. For instance, recent snRNA-Seq studies have revealed three transcriptionally distinct subclusters of human POMC neurons – variability that remained undetectable in our bulk datasets. Thus, IHC/LCM-Seq and single-cell approaches should be viewed as complementary methodologies, each offering unique advantages depending on the biological question.

### Functional implications of transcriptomic findings

Our study provides unprecedented insight into the molecular architecture of orexigenic AgRP and anorexigenic POMC neurons playing central roles in the regulation of energy homeostasis. The inclusion of KP neurons as a third cell type offers a valuable comparative perspective; while localized similarly in the arcuate nucleus, these neurons play a primary role in regulating the reproductive axis.^7^ With 14,000-16,000 transcripts detected per neuronal cell type, our dataset offers a comprehensive resource for dissecting the transcriptional programs that underlie the mediobasal hypothalamic control of energy balance and reproduction.

#### TFs reflecting shared and unique regulatory features and developmental origins of arcuate neuron phenotypes

Despite their distinct functional roles, AgRP, POMC, and KP neurons exhibited substantial overlaps in their TF profiles, similarities potentially reflecting shared developmental origins^13^ and/or common microenvironmental influences; among the 902 detected TFs, only 50 were differentially expressed across cell types.

TFs typical of AgRP neurons included *OTP, FOXO1*, and the glucocorticoid-responsive TFs *NR3C1*^14^*, KLF9*^15^, and *ZBTB16*^16^. Most TFs detected in POMC cells were not unique, as many were also expressed in AgRP or KP neurons. The presence of sex steroid receptors (*PGR*, *AR*, *ESR1*) provides a potential mechanistic link between gonadal hormone signaling and the hypothalamic regulation of metabolic functions^17,18^; not surprisingly, these receptors are expressed at the highest levels in KP neurons, consistent with the crucial involvement of this cell type in negative sex steroid feedback to the reproductive axis.^7^

#### Distinct transcriptomic signatures of AgRP, POMC, and KP neurons

Using differential expression analysis, we detected 1,210 transcripts exhibiting significant differences in expression across the three neuronal populations. *CARTPT* was expressed in AgRP neurons but not in POMC cells, representing a notable species difference from rodents that has already been reported previously.^19^ High levels of this mRNA were also detected in KP neurons, consistent with our laboratory’s earlier immunohistochemical observations.^20^ Additional neuropeptide mRNAs with enriched expression in AgRP neurons included *NPY, TAC1, SST, GHRH, GAL, CRH*, *VGF* and *UTS2.* Some of the encoded peptides may play thus far underappreciated roles in human orexigenic circuitry. *ADCYAP1* and *VIP* represented two neuropeptides expressed predominantly in POMC neurons and only at relatively low levels. Further transcripts typical of the POMC cell type – including *PENK*, *CBLN1*, *CBLN4*, *NXPH1*, and *PDYN* – were even more prominent in KP neurons, whereas *TAC3*, encoding neurokinin B, was exclusively expressed in KP neurons.

#### Genes associated with metabolic disorders

Somewhat unexpectedly, of the 2,493 transcripts associated with metabolic traits in the GWAS Catalog^10^, the majority (1,990) showed very similar expression across all three neuronal populations. Only 503 exhibited twofold or greater enrichment in a specific cell type, suggesting that many GWAS-linked genes may exert their effects across multiple hypothalamic pathways. Receptor genes from this database also tended to be expressed in multiple cell types, reinforcing the complexity of hypothalamic control of energy homeostasis.

#### Receptor transcriptomes hide novel therapeutic opportunities

A key objective of this study was to characterize receptor expression patterns with a focus on AgRP and POMC neurons to identify potential targets for metabolic therapies. In AgRP neurons, *ACVR1C* (ALK7) showed the highest expression. Genetic loss of this receptor or its natural ligand activin B in mice reduces the number of Npy-expressing cells, decreases *Npy* and *AgRP* mRNA levels, and diminishes NPY/AgRP projections to the MPOA, thereby supporting a trophic role for Alk7 in maintaining orexigenic circuitry.^21^ AgRP neurons highly expressed metabolic hormone receptors, including *GHSR* (for ghrelin), *INSR* (for insulin), and several cytokine receptors acting through the MyD88-dependent NF-κB and MAPK signaling cascade (*IL18R1*^22^, *IL1R1*^23^) and other receptors using JAK-STAT signaling (*GHR*^24^ and *LEPR*^25^*, IL6ST* and *IL13RA1*^26^). These receptors indicate that AGRP neurons serve as a key cell type of immune-metabolic integration. The receptor mRNA for PACAP, *ADCYAP1R1*, was highly expressed across all cell types. PACAP may exert a dual anorexigenic effect, analogous to leptin, by inhibiting AgRP neurons and activating POMC neurons within the arcuate nucleus, thereby promoting melanocortin signaling to downstream targets such as the paraventricular hypothalamus.^27^ The relaxin receptor *RXFP1* was also abundant in AgRP cells, while POMC neurons expressed *RXFP2.* In pregnant women, relaxin rises markedly, which promotes reproductive tissue remodeling, cardiovascular adaptation, and fluid homeostasis.^28^ Its action on AgRP neurons may align reproductive status with energy availability to support fetal development. Studies in rats found reduced food intake following peripheral or central administration of relaxin.^29^ The occurrence of the V1a vasopressin receptor *AVPR1A* in AgRP neurons may link hydration or stress state to feeding drive.^30^ The angiotensin II receptor *AGTR1* we detected in AgRP neurons may play an important role in resting metabolic rate adaptation, a mechanism which likely contributes to failed weight management during obesity.^31^ The interesting presence of the three olfactory receptors *OR52N5*, *OR52N1*, and *OR52H1* in AgRP neurons suggests potential responsiveness to yet-unidentified circulating metabolites or microbiota-derived compounds.

POMC neurons abundantly expressed *PRLR* and prolactin signaling on this receptor through the JAK-STAT pathway (as *IL6ST*^32^*, LEPR* and *IL13RA1*) may contribute to the coupling of reproductive status with metabolic regulation. The POMC cell type was also enriched in *CALCR* mRNA, which mediates responses to amylin and calcitonin – peptides known to promote satiety and to possibly act synergistically with leptin.^33^ The CGRP neuropeptide derived equally from the calcitonin gene (*CALCA)*, signals mainly through *CALCRL;* this receptor was found in the AgRP cell type. The serotonin receptor *HTR2C* was enriched in arcuate POMC neurons which may explain the anorexigenic actions of serotonin and the selective 5-HT2C agonists lorcaserin^34^ via modulating melanocortin output. Our results also shed light to some important species differences and limitations of rodent models. For example, while serotonin inhibits AgRP neurons primarily through *Htr1b* in rodents,^35^ human AgRP neurons did not express *HTR1B* but abundantly contained the *HTR2A* transcript. POMC neurons were highly enriched in *CNR1* (CB1 receptor). The endocannabinoid responsiveness of these neurons^36^ was proposed to enhance appetite via modulating glutamatergic signaling to the paraventricular nucleus.^37^ Orexin A acts via OX-1R to stimulate the endocannabinoid signaling in rodent POMC cells,^38^ whereas OX-2R receptors (encoded by *HCRTR2*) may exert a similar effect on POMC neurons in the human, as indicated by our results. Intact histamine signaling is necessary for proper feeding rhythm and optimal leptin signaling. Expression of *GLP1R* supports the hypothesized central mechanism of appetite suppression by semaglutide – a GLP-1 receptor agonist used in obesity and diabetes treatment.^39,40^ Our transcriptomic analysis confirmed the recent observation that the expression level of *GLP1R* in human POMC neurons is relatively low compared with mice.^4^ Although at much lower levels, *GLP1R* was also present in AgRP neurons. This finding corroborates recent observations on mice where the orexigenic cell group not only expresses GLP-1R protein but also receives input from axons that contain this receptor.^41^ The presence of μ-opioid (*OPRM1*), nociceptin/NOP (*OPRL1*), and κ-opioid (*OPRK1*) receptors in POMC neurons suggests mechanisms whereby opioid peptides may modulate appetite and hedonic feeding.^42–44^

Orphan GPCRs, including *GPR34, GPR27* (SREB1), and *GPR3* enriched in AgRP neurons, and *GPR101, GPR176*, *GPR83, GPR149*, and *GPR85* (SREB2) expressed in POMC neurons, and *GPR107, GPR75, GPRC5B, GPR158, GPR162, GPR135, GPR19, GPR137C, GPR137B, GPR61,* and *GPR6* present in both neuron populations, may represent particularly promising candidates for pharmacological modulation of hunger and satiety circuits.

Overall, comprehensive receptor profiling of human AgRP and POMC neurons in this study unveiled multiple receptors that play previously unrecognized roles in the hypothalamic regulation of energy homeostasis and offer drug targets for future metabolic therapies.

## Acknowledgments

This work was supported by grants to Project no. RRF-2.3.1-21-2022-00011, titled National Laboratory of Translational Neuroscience implemented with the support provided by the Recovery and Resilience Facility of the European Union within the framework of Programme Széchenyi Plan Plus, the National Research, Development and Innovation Office (K138137 to E.H. and PD134837 to K.S). We would like to thank Drs. C. Fekete and W.S. Dhillo for kind donation of the POMC and KP antibodies, respectively.

## Author contributions

**Conceptualization:** S.T., K.S., B.G., É.R., M.S., E.H.

**Investigation:** S.T., K.S., B.G., É.R., M.S., S.P., G.R., A.M., E.H.

**Writing-editing:** S.T., K.S., E.H.

**Methodology:** S.T., K.S., B.G., É.R., M.S., S.P., G.R, A.M., E.H.

**Funding acquisition and supervision:** K.S., E.H.

## Supplemental information titles and legends

**Table S1**: RIN and RNA yield values for **Figure 1B**, RIN values and ANOVA statistics for **Figure 1D**

**Table S2**: Whole transcriptome profile of hypothalamic AgRP, POMC and KP neurons including differential expression data

**Table S3**: Gene expression profile of transcription factors

**Table S4**: Gene expression profile of neuropeptides and granins

**Table S5**: Selected, metabolism-related mapped traits from the GWAS Catalog

**Table S6**: Transcripts filtered for GWAS association by mean TPM ≥ 10 and at least 5 reads in each sample of the same neuronal phenotype

**Table S7**: Transcripts of ‘MAPPED_GENEs’ expressed in AgRP, POMC or KP neurons that were associated with metabolic disorders in the GWAS Catalog

**Table S8**: Receptor expression profiles of AgRP, POMC and KP neurons including GWAS trait associations

## Methods

### Experimental model and subject details

#### Human subjects

Histological samples from adult human brains (N = 8) were obtained during autopsies conducted at the Department of Pathology and Experimental Cancer Research, Semmelweis University, Budapest, Hungary. The specimens were derived from one 53-year-old female and seven male individuals aged 28, 33, 38, 46, 52, 53, and 56 years, all with no documented neurological disorders. Tissue collection was performed 12 to 24 hours *postmortem*. Ethical approval was granted by the Regional and Institutional Committee of Science and Research Ethics of Semmelweis University (SE-TUKEB 251/2016), in compliance with the Declaration of Helsinki and relevant Hungarian legislation (Act CLIV of 1997 and Decree 18/1998 [XII.27.] of the Ministry of Health), which does not mandate prior written consent from the deceased.

### Method details

#### Experimental procedures

For all experiments, reagents were of molecular biology grade. MQ water and PBS were pretreated overnight with diethylpyrocarbonate (DEPC; Thermo Scientific; 1 mL/L) and subsequently autoclaved. All other solutions were prepared using DEPC-treated and autoclaved MQ water. Reagents and buffers were maintained under RNase-free, sterile conditions with or without the additional use of the RNase inhibitors. Work surfaces were cleaned with RNaseZAP (Merck KGaA, Darmstadt, Germany).

#### Optimization of the IHC/LCM-Seq method for human brain tissues

##### Immersion fixation and section preparation

To optimize the IHC/LCM-Seq protocol for brain samples gained from autopsies, test experiments were performed using 4-mm-thick tissue blocks (n = 9) dissected 20 h *postmortem* from the frontal cortex of a 53-year-old female subject. Samples were immersion-fixed for 24, 48, or 72 hours (n = 3 per fixation time) in freshly prepared ice-cold 4% formaldehyde (FA) in 0.1 M phosphate-buffered saline (PBS; pH 7.4). Following fixation, tissues were infiltrated with ice-cold sterile 20% sucrose in PBS for 24 hours, snap-frozen on powdered dry ice, and stored at –80 °C until sectioning. The fixative and sucrose solutions were supplemented with 0.05% aluminon (triammonium salt of aurintricarboxylic acid, ATA; Merck) to enhance RNA preservation.^6,8^ Prior to sectioning, the white matter was trimmed from each tissue block. Free-floating 20-µm-thick sections were prepared using a Leica freezing microtome (Leica Biosystems, Wetzlar, Germany) (**Fig. 1A**).

##### Assessing the impact of immersion fixation on RNA quality and recovery

RNA was extracted from individual cortical sections fixed for 24, 48, or 72 hours using the PureLink FFPE RNA Isolation Kit (Thermo Scientific, Waltham, MA, USA). For each fixation time (n = 3), RNA yield – normalized to the section area (mm^2^) – and RNA Integrity Number (RIN) were determined using the Agilent 2100 Bioanalyzer system with Eukaryotic Total RNA Pico Chips and the 2100 Expert software (Agilent, Santa Clara, CA, USA). Results were presented as mean ± SEM from triplicate measurements (**Table S1**).

##### Tissue archiving and optimization of RNA preservation during antigen retrieval

Most cortical sections were stored for one month at –20 °C in a modified cryoprotectant solution (30% ethylene glycol, 25% glycerol, 0.05 M phosphate buffer; pH 7.4) supplemented with 0.05% aluminon (ATA). We have previously demonstrated that this storage method preserves RNA integrity for three years or longer.^6^ Optimization of section pretreatments and the IHC protocol used the three tissue blocks that were immersion-fixed for 72 hours. Sections retrieved from the antifreeze medium were rinsed in sterile PBS and subjected to a 30-minute heat-induced epitope retrieval step in 0.1 M citrate buffer (pH 6.0, 80 °C), either without additives or in the presence of the RNase inhibitors ATA (0.05%) or polyvinyl sulfonic acid (PVSA)^45^ used at 1%, 3%, or 10%. Following brief PBS washes, total RNA was extracted, and RNA integrity was assessed using the Agilent 2100 Bioanalyzer system. Results were presented as mean ± SEM (**Table S1**) of three measurements on independent samples. Differences between treatment groups – with and without antigen retrieval – were assessed by one-way ANOVA followed by Tukey’s *post hoc* tests.

##### Optimization of the RNA-preserving immunohistochemistry (IHC) protocol

Antigen retrieval was performed in the presence of 10% polyvinyl sulfonic acid (PVSA) which efficiently protected RNA in the above experiment. Subsequently, sections were processed for immunohistochemical detection of cortical parvalbumin-expressing neurons at either RT or 4 °C, with or without the use of RNase inhibitors. To inhibit RNases, all antibody and buffer solutions (except the peroxidase developer) contained either 0.05% aluminon (ATA) or 2% PVSA. Sections were pretreated with 1% H_2_O_2_ and 0.5% Triton X-100 in PBS for 20 minutes, followed by incubation with a commercially available affinity-purified goat polyclonal antibody against human parvalbumin (#EB06776; Everest Biotech; 1:30,000) for either 40 hours at 4 °C or overnight at RT. The primary antibody was reacted with Biotin-SP-AffiniPure anti-goat IgG secondary antibody (Jackson ImmunoResearch; 1:500; overnight at 4 °C or 2 hours at RT), followed by the ABC Elite reagent (Vector Laboratories, Burlingame, CA, USA; 1:1,000; 1 hour at RT) in PBS. The peroxidase reaction was visualized at RT using a developer containing 10 mg diaminobenzidine, 30 mg nickel-ammonium sulfate, and 0.003% H_2_O_2_ in 20 ml of 0.05 M Tris-HCl buffer (pH 8.0). RNA was extracted from pre-IHC sections (n = 3) and from post-IHC sections (n = 3 per treatment group), and RNA integrity was assessed using the Agilent 2100 Bioanalyzer System (Table S1). Differences in RIN values among the post-IHC groups were analyzed using one-way ANOVA followed by Tukey’s *post hoc* test.

For qualitative analysis of the immunohistochemical labeling, the immunostained sections were mounted onto microscope slides from Elvanol, air-dried for 30 minutes, then sequentially dehydrated in 70%, 95%, and 100% ethanol (5 minutes each), followed by clearing in xylene (2 × 5 minutes). Coverslipping was performed using DPX mounting medium (Merck, Darmstadt, Germany) for light microscopic analysis and imaging. Digital images were adjusted for brightness and contrast using Adobe Photoshop (Adobe Systems), and final composite figures were assembled in Adobe Illustrator (Adobe Systems).

#### IHC/LCM-Seq studies of hypothalamic AgRP, POMC and KP neurons

##### Immersion fixation of hypothalamic samples

Four-mm-thick hypothalamic slices from seven male individuals (**Fig. 2C**) were immersion-fixed in ice-cold 4% FA in 0.1 M phosphate-buffered saline (PBS; pH 7.4) for 72 hours, followed by infiltration in 20% sucrose (24 h). Both solutions were supplemented with 0.05% aluminon (ATA). Free-floating 20-µm-thick coronal sections were prepared from the mediobasal hypothalamus (atlas plates 24-27)^46,47^ using a freezing microtome and stored in modified cryoprotectant solution containing 0.05% ATA, as described for cortical samples.

##### RNA quality testing

Total RNA was extracted from a hypothalamic section per case using the PureLink FFPE RNA Isolation Kit (Thermo Scientific) its integrity was assessed with the Agilent 2100 Bioanalyzer system. Samples used in subsequent IHC/LCM-Seq experiments exhibited RNA Integrity Number (RIN) values ranging from 3.1 to 6.7 (**Fig. 2C**).

##### Immunohistochemical detection of AgRP, POMC and KP neurons

The RNA-preserving IHC protocol optimized for cortical parvalbumin labeling was adapted to detect hypothalamic AgRP, POMC and KP neurons. All antibody and buffer solutions were supplemented with 2% PVSA, except the epitope retrieval solution, which contained 10% PVSA, and the peroxidase developer, which was free of RNase inhibitors.

Mediobasal hypothalamic sections (every 12^th^) were removed from the cryoprotectant solution, rinsed in ice-cold PBS (3 × 5 min), subjected to antigen retrieval in 0.1 M citrate buffer (pH 6.0; 80 °C; 30 min), then incubated at 4 °C for 20 minutes in PBS containing 1% H_2_O_2_ and 0.5% Triton X-100. The following primary antibodies were applied for 40 hours at 4 °C: Goat anti-AgRP (EB06725; Everest Biotech; 1:10,000); Sheep anti-POMC (#41; gift from Dr. Csaba Fekete; 1:30,000)^48^; Sheep anti-KP (GQ2; gift from Dr. Waljit S. Dhillo; 1:50,000)^49^. After PBS washes, sections were incubated for 2 hours at RT with species-specific biotinylated secondary antibodies raised in donkeys (Jackson ImmunoResear ch; 1:500), followed by a 1-hour incubation with the ABC Elite reagent (Vector Laboratories, Burlingame, CA, USA; 1:1,000) in PBS. Immunoreactivity was visualized using Ni-DAB chromogen as described for cortical parvalbumin labeling.

##### Selection and exclusion criteria of immunostained sections for RNA-Seq studies

Due to variability in staining quality, not all seven hypothalamic specimens were included in all downstream analyses. Transcriptome profiling was conducted using: 3 samples for AgRP neurons, 2 samples for POMC neurons and 3 samples for KP neurons

##### Laser capture microdissection of immunostained neurons

Immunolabeled sections were mounted onto PEN membrane glass slides (Membrane Slide 1.0 PEN, Carl Zeiss) from Elvanol containing 2% PVSA, air-dried, and processed for LCM via sequential dehydration: 70% ethanol (60 s), 96% ethanol (60 s), 100% ethanol (2 × 60 s), and xylene (60 s). Slides were stored at –80 °C in slide mailers with silica gel desiccants unless processed immediately for LCM. Using a PALM Microbeam system (Carl Zeiss) equipped with a 40× objective lens and RoboSoftware, immunolabeled neurons were individually selected from the arcuate/infundibular nucleus and pressure-catapulted into adhesive caps (Adhesive Cap 500, Carl Zeiss). Approximately 850 neurons were collected per sample and stored at –80 °C in LCM caps until RNA extraction.

##### RNA extraction

Total RNA was isolated from pooled neurons using the Arcturus Paradise PLUS FFPE RNA Isolation Kit (Applied Biosystems, Waltham, MA, USA), according to the manufacturer’s protocol.

##### Library preparation and RNA Sequencing

Sequencing libraries were generated with the TruSeq Stranded Total RNA Library Preparation Gold kit (Illumina, San Diego, CA, USA), incorporating 16 PCR cycles for fragment enrichment.^6,50^ A pooled library mix (1 nM; 120 µl total volume) containing eight indexed samples was sequenced on an Illumina NextSeq 500 platform using the NextSeq 500/550 High Output v2.5 kit (75-cycle configuration).^11^

##### Bioinformatics

Low-quality reads and adapter sequences were removed using Trimmomatic (v0.39; parameters: LEADING:3, TRAILING:3, SLIDINGWINDOW:4:15, MINLEN:36)^52^ and Cutadapt (v4.6)^51^, respectively. Cleaned reads were aligned to the *Homo sapiens* GRCh38 Ensembl reference genome (release hs111) using the STAR aligner (v2.7.11b)^53^ with default parameters. Gene-level quantification was carried out using featureCounts (Subread v2.0.6).^54^ Transcriptome coverage was assessed based on the number of unique transcripts detected at two read count thresholds: ≥1 and ≥5 reads. Metrics were calculated and reported as the mean ± standard error of the mean (SEM) across biological replicates. Subsequent analysis was conducted in R (v4.5.0).^55^ TPM values were computed from read counts normalized to gene lengths obtained via FeatureCounts, using gene models defined in the corresponding Ensembl GTF annotation file. For differential expression analysis, raw gene counts were normalized and analyzed using the DESeq2 package (v1.44.0).^56^ The following R packages were utilized for data transformation, analysis, and visualization: tidyverse (v2.0.0)^57^, openxlsx (v4.2.8)^58^, ggplot2 (v3.5.2)^59^, ggfortify (v0.4.17)^60^, cluster (v2.1.8.1)^61^, pheatmap (v1.0.12)^62^, RColorBrewer (v1.1-3)^63^, ggrepel (v0.9.6)^64^, EnhancedVolcano (v1.22.0)^65^, gridExtra (v2.3)^66^, ggpubr (v0.6.0)^67^, eulerr (v7.0.2)^68^.

##### Functional classification

Transcripts were assigned to functional categories using the KEGG BRITE database^69^ (https://www.genome.jp/kegg/brite.html). Transcriptional regulators were identified based on the KEGG ‘Transcription Factors’ category (ko03000) (see **Table S3**). Receptor classes included G protein-coupled receptors (ko04030), cytokine receptors (ko04050), pattern recognition receptors (ko04054), and nuclear receptors (ko03310). This functional category was extended to include ACVR1C and INSR from the receptor serine/threonine kinases and receptor tyrosine kinases categories (ko01001+K08873), respectively. Receptor genes were considered expressed in a given neuronal phenotype if their mean TPM value was ≥10 and they were supported by ≥5 reads in all biological replicates of that group (**Table S8**). Neuropeptides were classified based on the neuropeptides.nl database^9^ (**Table S4**). A comprehensive list of functional classifications is provided in **Tables S3, S4, S8**.

##### Populational enrichment of gene expression

Gene expression was considered enriched in a specific neuronal population if the transcript exhibited at least a twofold increase in normalized read counts (log_2_ fold change > 1) relative to the other populations. When two neuronal types each showed ≥2-fold higher expression compared to the third, the gene was classified as co-enriched in those two populations.

##### GWAS associations

Trait– and disease-associated SNPs were retrieved from the GWAS Catalog^10^ (https://www.ebi.ac.uk/gwas/). Traits and diseases relevant to malnutrition were manually curated to match the 14 mapped GWAS traits presented in **Figure 5** and **Table S5**. Trait categories included all descendant terms, except for ‘Body weight’, where child terms were excluded to maintain specificity. For comparative analysis, only intragenic SNPs reaching genome-wide significance (p < 5 × 10^-8^) were retained. The resulting gene list was then cross-referenced with neuronal population-specific enrichment data (**Tables S6, S7**).

##### Data availability

RNA-Seq files for AgRP and POMC neurons are available in BioProject with the accession number PRJNA1305226. (Release date 01/01/2026). KP sequencing data have been deposited under PRJNA1305334 (Release date 01/01/2026).

##### Code availability

Scripts are available at https://github.com/goczbalazs/PRJNA1305226_1305334.

##### Quantification and statistical analysis

Statistical analyses for **Figure 1D** were conducted using one-way ANOVA followed by Tukey’s HSD *post hoc* tests, implemented via the aov() and TukeyHSD() functions in R. Normality was evaluated with the Shapiro–Wilk test, and homogeneity of variances was assessed using Levene’s test (Brown–Forsythe; median-centered) – via the shapiro.test() and leveneTest() functions. Significance was set at p < 0.05. Full statistical results are presented in **Table S1.**

Differential expression analysis for RNA-Seq data was performed with the DESeq2 package. Genes with Benjamini-Hochberg adjusted p-values < 0.05 were considered differentially expressed. Detailed results are provided in **Table S2**.

Significant SNP-trait associations from GWAS were defined by p < 5×10^-8^.^70,71^ The annotation and filtering of SNPs by intragenic localization and trait mapping are described above under GWAS Associations.

Codes are available at https://github.com/goczbalazs/PRJNA1305226_1305334

### Resource availability

#### Lead contact

Further information and requests for resources and reagents should be directed to and will be fulfilled by the lead contact, Erik Hrabovszky.

#### Materials availability

This study did not generate new unique reagents.

